# A Retro-Inverso Peptide that Inhibits Formation and Disrupts Preformed α-Helical Amyloid and Staphylococcus aureus Biofilms

**DOI:** 10.64898/2026.04.24.720651

**Authors:** Abhisek Kar, Atlanta Borah, Vishal Malik, Anand Srivastava, Kalyaneswar Mandal

**Affiliations:** Tata Institute of Fundamental Research Hyderabad, 36/p Gopanpally, Hyderabad, Telangana – 500046, India; National Institute of Animal Biotechnology (NIAB), Gowlidoddy, Gachibowli, Hyderabad, Telangana - 500032, India

**Keywords:** Staphylococcus aureus, Phenol-soluble modulins, amyloid, fibrils, biofilm, Retro-inverso peptide

## Abstract

Combatting biofilm-associated methicillin-resistant Staphylococcus aureus (MRSA) infections remains a formidable challenge, largely due to bacteria’s heightened resistance to antibiotics and evasion of the host’s immune responses. Inhibiting biofilm formation or promoting biofilm dissolution is believed to disrupt the protective matrices and expose bacterial cells to immune clearance or antimicrobial treatments. Cross-α helical amyloid fibrils of phenol-soluble modulins are the key structural components of the MRSA biofilm. In this study, we present a novel retro-inverso peptide that successfully inhibits cross-α amyloid formation, induces disassembly of pre-formed fibrils of the biofilm-forming modulin peptide, and efficiently disperses MRSA biofilm biomass in a dose-dependent manner. The biophysical studies of the interaction of the designed peptide and modulins have suggested a mechanism for the biofilm disruption. The findings highlight the potential of the proteolytically resistant mirror-image peptide as a promising therapeutic agent for the treatment of biofilm-associated MRSA infections.

## Introduction

The self-aggregation of proteins and peptides forms amyloids into highly ordered fibrillar structures.^1-4^ The amyloid form of proteins and peptides are responsible for many human degenerative pathologies.^1,5,6^ The high thermodynamic stability of such protein and peptide aggregates resists disruption of amyloids by proteases and detergents.^7-9^ Such amyloid aggregates are the primary component of the biofilm of several bacterial species.^10-13^ The biofilm protects bacterial cells from the host immune responses, antibiotics, and disinfectants.^14-18^ *Staphylococcus aureus* (*S. aureus*), a gram-positive bacterium, is frequently found in the nose, respiratory tract, and on the skin, and causes infections such as endocarditis, necrotizing pneumonia, and septic shock.^19^ The major constituents of *S. aureus* biofilm amyloid fibrils are small amphipathic phenol soluble modulins (PSMs).^20-26^ In previous studies, soluble PSMs have been shown to facilitate biofilm dissolution.^[11]^ In contrast, another study found that the PSM peptides exist as fibrils that not only self-organize but also support the structural integrity of the biofilm.^35^ Thus, amyloid-like aggregation regulates the dual and opposing roles played by PSMs.

The amphipathic nature of several PSMs enables them to function as biological detergents, facilitating their ability to permeate cell membranes.^36^ Among all the PSM family peptides, PSMα3, an α-type modulin from *S. aureus,* containing 22 amino acids, exhibits the highest cytolytic activity.^31,37^ While the biological activity of these PSMs is well addressed, the structural details that govern their interactions with the biological systems have yet to be fully established. Recently, in a notable study, Landau and co-workers reported the formation of a cross-α amyloid-like fibrillar structure from 22-residue PSMα3 peptide.^38^ Alpha-helical cross-α amyloid-like fibrillar structure is unique in the sense it has not been seen in other amyloid structures, which are usually beta-sheet aggregates. They hypothesized that this cross-α amyloid-like fibrillar structure is possibly responsible for the greater cytotoxicity of PSMα3. However, the precise structural identity of the molecular assembly that contributes to the activity of PSMα3 has still remained elusive. Moreover, the mechanisms that influence the ability of PSMs to transition from a soluble form to a fibril state are currently unknown, and it is unclear how this affects biofilm development or degradation. It is believed that PSMs in a sessile biofilm remain as inert fibrils until cellular conditions allow them to dissociate, manifesting biofilm’s antimicrobial activity, virulence, or degradation.^35^

Despite the collective efforts of many research groups, combatting infections triggered by *S. aureus* remains a formidable treatment challenge, primarily due to the bacteria’s heightened resistance to antimicrobial treatments and the host’s immune responses. This resistance originates in the *S. aureus* biofilm. We presumed if PSMs’ insoluble fibrillar state is an integral part of biofilms, preventing their formation or promoting the dissolution of the pre-formed fibrils would be an effective way to disperse the biofilm matrices and expose the cell to the host immune responses or antibacterial treatments. This approach goes beyond conventional inhibitors by actively disrupting mature amyloid structures without inducing resistance. The mechanism avoids selective pressure and enhances combinatorial efficacy with antibiotics by weakening the biofilm rather than killing bacteria directly. We embarked on identifying peptide-based inhibitors that can impede the development of the extracellular polymeric substance (EPS) matrix and, more importantly, disassemble the fully formed EPS matrix within *S. aureus* biofilms. In pursuit of this objective, we systematically investigated the effect of a retro-inverso PSMα3 (RI-PSMα3) on the cross-α amyloid assembly of L-PSMα3 (native PSMα3), facilitating *S. aureus* biofilm dissolution.

In retro-inverso peptide, the amino acid sequence is reversed (retro) and each amino acid is replaced with its enantiomer (inverse) (*see Figure 1A*). Despite the reversed sequence and inversed chirality, retro-inverso peptides, especially those with an α-helical structure, can interact with the same receptor as the parent peptide because the side chain topology of the retro-inverso sequence can mimic the original peptide’s conformation.^46^ Additionally, retro-inverso peptide has the advantage of increased resistance to proteolysis due to the presence of D-amino acids.^47^

**Figure 1.**
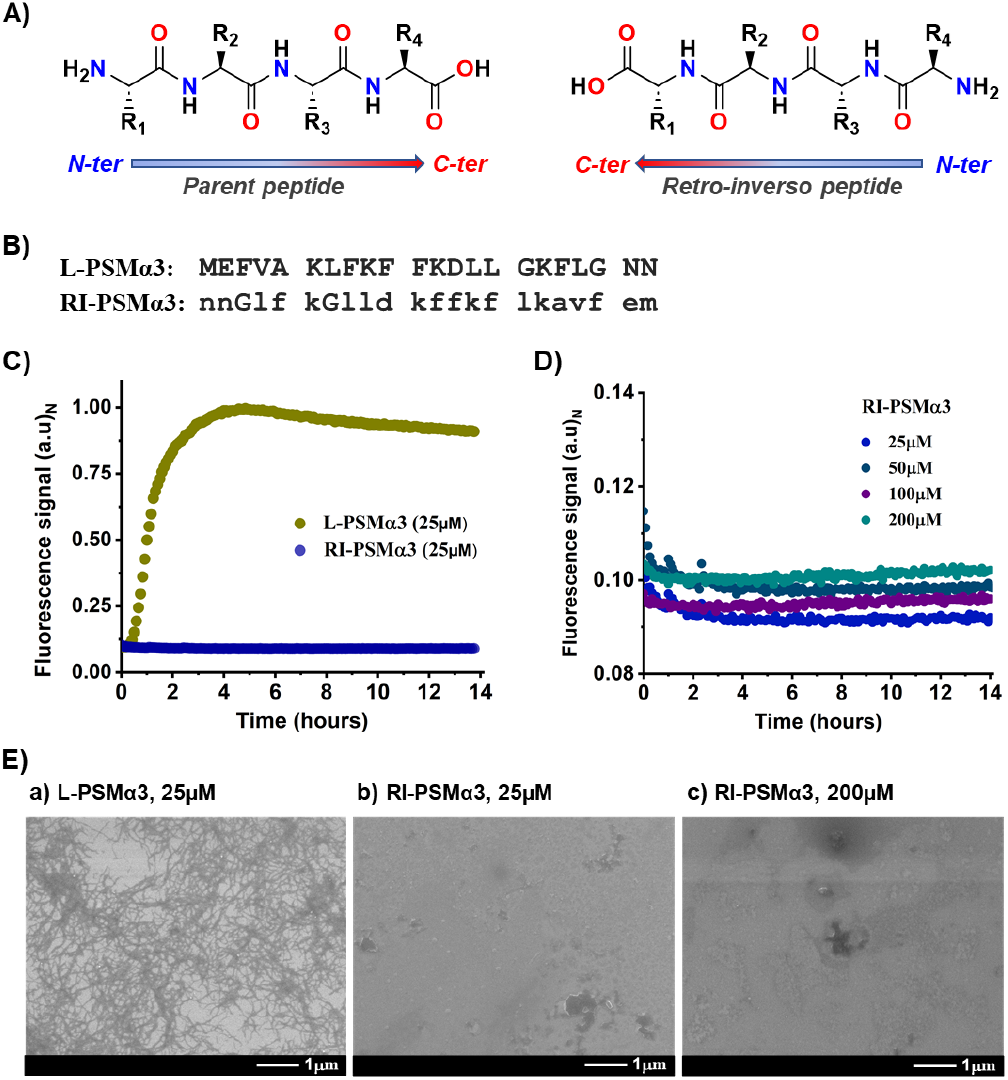
Aggregation of the PSMα3 peptides. (A) Topological relationship shown for a model peptide and its fully retro-inverso analog. B) Sequence of PSMα3 and RI-PSMα3 peptides. C) Aggregation of L-PSMα3 and RI-PSMα3 were monitored by ThT fluorescence assay. The concentration of each peptide was 25 μM. D) Aggregation of the RI-PSMα3 peptide at increased concentrations (25 μM, 50 μM, 100 μM, and 200 μM) as monitored by the same ThT fluorescence assay. E) Representative SEM images of the a) L-PSMα3 (25 μM), b) RI-PSMα3 (25 μM), and c) RI-PSMα3 (200 μM) after 14 hours of incubation.

## Results and Discussion

We set out to understand the property of the retro-inverso (RI) sequence of the cross-α amyloid-forming peptide L-PSMα3, the most lytic member of the PSM family peptides of *S. aureus*. We prepared L-PSMα3 and RI-PSMα3 peptides as well as their Cysteine-containing analogs using Fmoc chemistry solid phase peptide synthesis (*see SI Section 1.2 for details*). The crude peptides were purified using reverse-phase HPLC (RP-HPLC), as shown in *SI Table S1*. The purified peptides were then subjected to a pre-treatment protocol with a TFA-HFIP (1:1) mixture (*see SI Section 1.3)*. This pretreatment procedure was designed to effectively dissect all aggregates present in the sample of interest.^48,49^

### RI-PSMα3 peptide does not form amyloid

The time dependent thioflavin-T (ThT) aggregation assay represents a well-established method for demonstrating the formation of amyloid aggregates under the experimental conditions used.^50,51^ The ThT assays with the L-PSMα3 and RI-PSMα3 have been detailed in *SI Section 1.5*. In the ThT fluorescence assay for the L-PSMα3 peptide at 25 μM concentration, we observed a very short initial lag phase, a rapid growth phase followed by the plateau phase, indicative of equilibrium between fibril formation and fibril dissociation or fragmentation (*Figure 1C*). This positive outcome is consistent with the previous studies.^38,45^ Interestingly, and to our surprise, at an equal concentration of RI-PSMα3, we did not observe any change in fluorescence signal, suggesting that, unlike L-PSMα3 peptide, RI-PSMα3 does not form cross-α type amyloid fibril (*Figure 1C*). The visualization of the samples using scanning electron microscopy (SEM) after 14 hours of incubation (*see SI Section 1.6*) revealed the presence of the amyloid formation for the L-PSMα3 peptide (*Figure 1E-a*), but not the RI-PSMα3 peptide (*Figure 1E-b*). Importantly, RI-PSMα3 did not exhibit any fibril formation even at a concentration as high as 200 μM (*Figure 1D & 1E-c*).

The formation of the cross-alpha amyloid structures in the L-PSMα3 peptide is driven by the thermodynamically favorable stacking of peptides in a parallel orientation, leading to the assembly of higher-order oligomers followed by nucleation and the eventual formation of insoluble fibrils.^38,52^ We hypothesized that due to its similar sidechain orientation, the RI-PSMα3 would interact with the native PSMα3 upon co-incubation, thereby preventing nucleation and fibril formation; and that sufficiently strong interactions might also disrupt preformed fibrils.

### RI-PSMα3 prevents L-PSMα3 amyloid formation

To assess the potential inhibitory effect of RI-PSMα3 on amyloid formation of the L-PSMα3 peptide, we conducted ThT assays in the presence of RI-PSMα3 (*see SI Section 1.7*). When solutions containing L-PSMα3 were incubated with an equal proportion of RI-PSMα3 in 10 mM sodium phosphate buffer containing 150 mM NaCl at pH 7.4 in the presence of 200 μM of ThT for 14 hours, a complete suppression of the fluorescence signal was observed (*Figure 2A*). We observed similar results irrespective of the peptide concentration used (12.5 μM, 25 μM, or 50μM) in the assay (*Figure 2A*). These ThT assay results were further substantiated by SEM analysis, which visually confirmed the inhibition of amyloid formation in the presence of RI-PSMα3 (*Figure 2B*).

**Figure 2.**
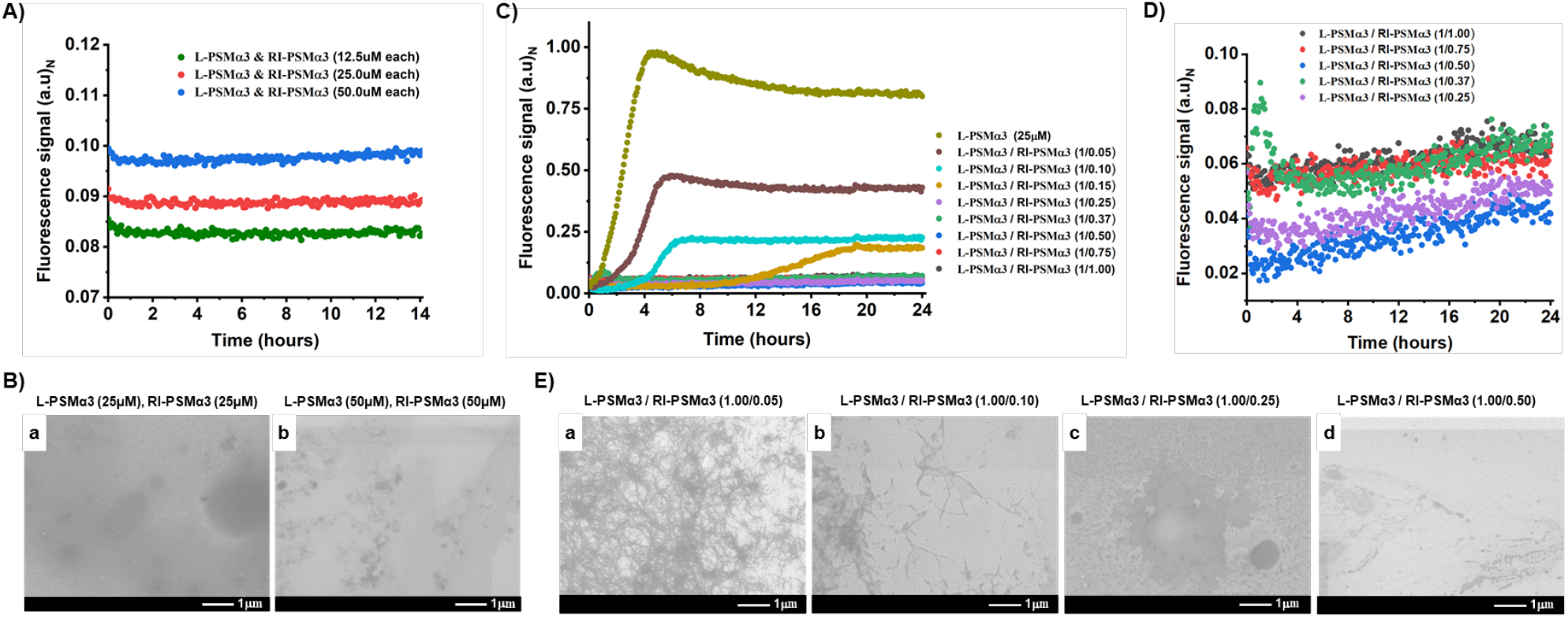
Inhibition of amyloid formation by L-PSMα3 peptide in presence of the RI-PSMα3 peptide. (A) The kinetics of L-PSMα3 amyloid formation in the presence of the RI-PSMα3, monitored by ThT fluorescence. Keeping the proportion of the L-PSMα3 and RI-PSMα3 peptides same, the concentration of each peptide varied from 12.5μM to 50μM. B) The representative SEM images of L-PSMα3 fibrils after 14 hours of incubation in the presence of an equimolecular amount of the RI-PSMα3 peptide. C) Dose-dependent PSMα3 amyloid formation in the presence of the RI-PSMα3, monitored by ThT fluorescence. The molar equivalent of the RI-PSMα3 peptide varied from 0.05 to 1.00 with respect to the L-PSMα3 peptide. D) The zoomed-in version of Figure C visualizes the complete suppression of the L-PSMα3 ThT intensity when the equivalent amount of the RI-PSMα3 peptide was 0.25 and above. E) The representative SEM images of L-PSMα3 fibrils after 24 hours of incubation in the absence or presence of the different equivalent amounts of RI-PSMα3 peptide.

Furthermore, we conducted co-incubation experiments in which L-PSMα3 peptide was combined with increasing concentration (equivalent amounts: 0.05, 0.10, 0.15, 0.25, 0.37, 0.50, 0.75, 1.00) of RI-PSMα3 peptide, while maintaining a constant L-PSMα3 concentration of 25μM (*see SI Section 1.8*). Relative to the untreated L-PSMα3, considered as 100% aggregation, the presence of RI-PSMα3 dramatically reduced the level of amyloid fibrils (*Figure 2C*). Even with a 0.25 molar equivalent of RI-PSMα3 peptide, no increase in the ThT fluorescence signal was observed during the 14-hour incubation period. The inhibition of L-PSMα3 fibril formation by RI-PSMα3 was found to be dose-dependent. The visualization of samples by SEM further underscores the dose-dependent reduction of amyloid formation in the presence of RI-PSMα3 (*Figure 2E a-d*). The above experiments comprehensively demonstrate the inhibitory efficacy of RI-PSMα3 on the cross-alpha amyloid formation of the L-PSMα3 peptide, highlighting the potential of RI-PSMα3 as a valuable peptidic agent for modulating amyloid formation by L-PSMα3 peptide.

### RI-PSMα3 effectively disassembles pre-formed L-PSMα3 amyloid fibrils

From a disease perspective, disassembling pre-existing amyloid aggregates and fibrils is highly desirable, as these structures contribute to pathogenesis. To assess the potential of the RI-PSMα3 peptide to disrupt pre-formed L-PSMα3 fibrils, we conducted a series of experiments to investigate its ability to disassemble these amyloid structures (*see SI Section 1.9*). Initially, we allowed the L-PSMα3 peptide to form aggregates. The progress of the fibril formation was monitored using the ThT assay. After 4.5 hours, when the ThT fluorescence intensity reached maximum (*Figure 3A*), varied concentrations of RI-PSMα3 peptide were introduced to the matured fibrils, followed by continuous monitoring of the ThT fluorescence intensity for the subsequent 11 hours. The results revealed a striking difference in the signal behavior. In the absence of RI-PSMα3, the ThT signal remained at the plateau, indicating the stability of the pre-formed L-PSMα3 fibrils. However, upon immediate addition of RI-PSMα3, we observed a drastic reduction of the ThT signal, indicative of the disruption of the pre-formed fibrils and oligomers in a dose-dependent manner (*Figure 3A*). Notably, the most significant reduction of the ThT signal, approximately 80% disaggregation, was observed when a 1.5-fold excess of RI-PSMα3 was introduced (*Figure 3B*), which was further supported by the SEM analysis (*Figure 3C*). In untreated preformed L-PSMα3 assemblies, a fibrillar network characteristic of amyloid presence was evident (*Figure 3C-a*). However, in the presence of RI-PSMα3, the fibril density gradually diminished (*Figure 3C b-d*). The magnitude of this effect varied depending on the molar equivalent of the RI-PSMα3 peptide used in the study. Intriguingly, no trace of fibrils was detectable in the presence of one equivalent of RI-PSMα3, suggesting an apparent transformation of the preformed fibrils into smaller soluble oligomers or monomers. These smaller species were not detectable by ThT assay, further emphasizing the disaggregation potential of RI-PSMα3.

**Figure 3.**
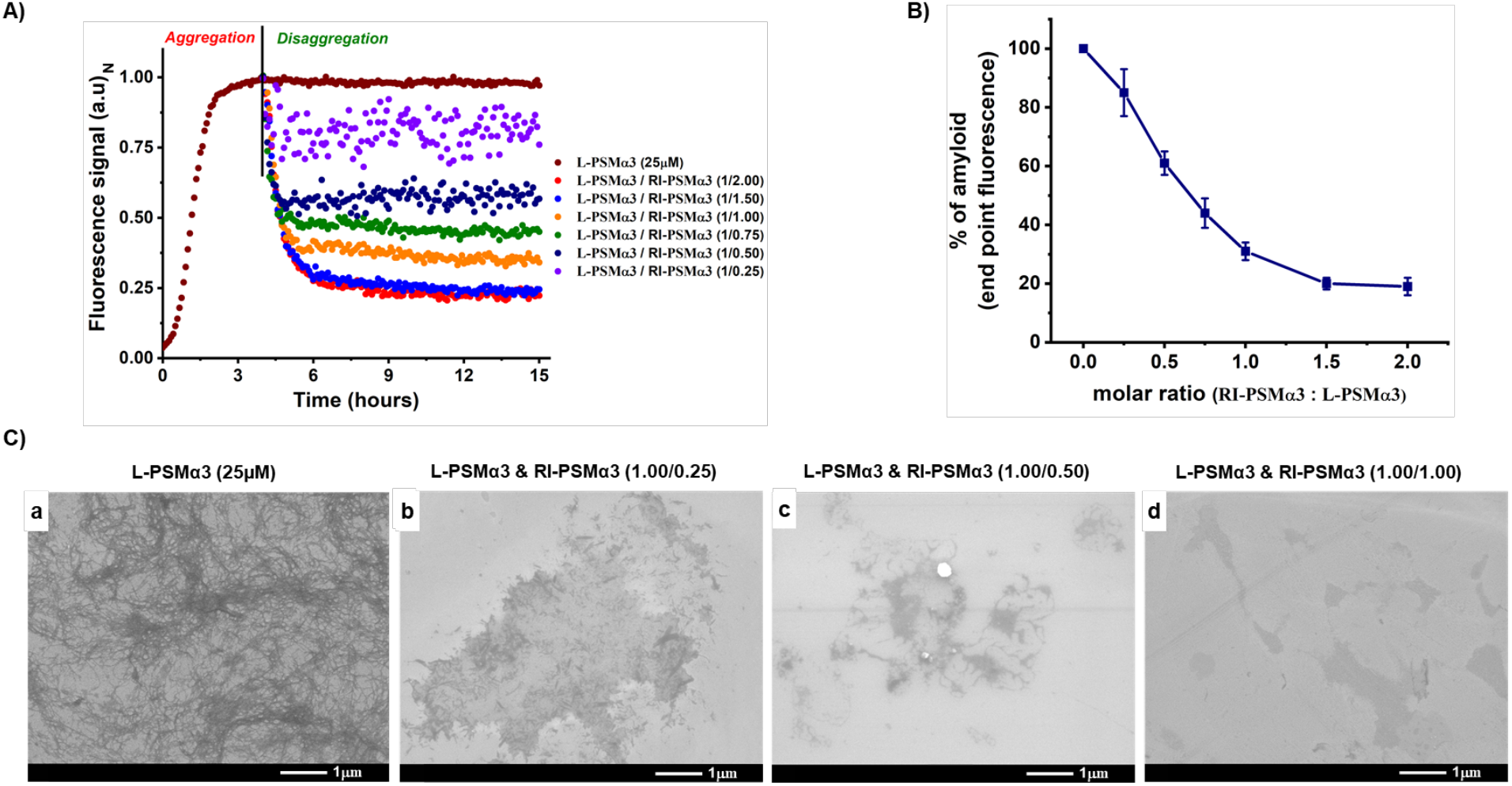
Dose-dependent disaggregation of L-PSMα3 amyloid by RI-PSMα3 peptide. (A) Dose-dependent ThT fluorescence measurement for the disaggregation of preformed fibrils of PSMα3 in the presence of RI-PSMα3. The molar equivalent of the RI-PSMα3 peptide was varied from 0.25 to 2.00 with respect to the L-PSMα3 peptide’s amount. B) Dose-dependent disaggregation of the preformed PSMα3 fibrils from end-point fluorescence data taken after 15 hours. C) The representative SEM images of pre-formed PSMα3 fibrils treated with different equivalents of RI-PSMα3 peptides.

These findings highlight the remarkable ability of the RI-PSMα3 peptide to effectively disassemble pre-formed amyloid fibrils, offering promising prospects for its therapeutic potential in mitigating the detrimental effects of PSMα3 associated amyloid-forming diseases. The observation of fibril transformation into soluble species underscores the potential of RI-PSMα3 in promoting the clearance of amyloid deposits.

### RI-PSMα3 binds with L-PSMα3 in an antiparallel fashion as determined by dynamic disulfide bond formation

To qualitatively decipher the mode of binding between L-PSMα3 and RI-PSMα3, we performed a dynamic disulfide bond formation reaction. This approach relies on the equilibrium distribution of a library of molecules containing disulfide bonds, which are under constant interconversion among themselves. Certain molecular species within the dynamic combinatorial library can be selectively stabilized or amplified under favorable conditions, depending on their thermodynamic properties influenced by interactions with other molecules.^53,54^ Dynamic covalent bond formation chemistry has been employed in various fields of chemistry, including supramolecular chemistry, drug discovery, and the study of biomolecular interactions.^53-58^ It is particularly useful for exploring and identifying molecules with unique properties or functions from a complex mixture of compounds.

Using dynamic combinatorial disulfide bond formation chemistry, we investigated the binding mode and affinity of the native L-PSMα3 and RI-PSMα3 peptides (*see SI Section 1.10*). Here, we hypothesized that the extent of disulfide bond formation between two peptides would reflect the extent of interaction between them, forming the thermodynamically stable homo- or hetero-dimer. To achieve this goal, we rationally designed and chemically synthesized peptides with cysteine residue either at the C-terminus or the N-terminus, enabling the use of dynamic disulfide bond formation chemistry with LC-MS as the read-out. Since all analogs share the same amino acid compositions, an additional (arbitrary) residue was introduced at either the N- or C-terminus of each peptide to distinguish them using mass-spectrometry (*see Table 4A*).

A schematic of the dynamic combinatorial disulfide bond formation chemistry is illustrated in Figure 4B. The experiment relied on the disulfide bond formation between two peptides under GSH/GSSG mediated folding conditions (20mM phosphate, 1mM GSH, 0.2mM GSSG, pH 8.0) followed by denaturation and reverse-phase high-performance liquid chromatography (RP-HPLC) coupled with online electron spray mass spectrometry (ESI-MS) analysis. The first three experiments in which individual peptides (*Figure 4_Table C, Exp1-Exp3*) were treated under redox folding conditions revealed a quantitative homo-dimer product, as expected, with traces of glutathione adduct formation (*Figure 4D*). In the next two experiments (*Figure 4_Table C, Exp4-Exp5*), we incubated equimolecular amounts of both peptides under redox folding conditions. When the C-terminal cysteine-containing peptide L-PSMα3(Cys) and RI-PSMα3(Cys) were incubated, the equilibrium product distribution followed statistical pattern (*Figure 4D, Expt. No. 4*). However, when the C-terminal cysteine-containing L-PSMα3(Cys) peptide was incubated with the N-terminal cysteine-containing (Cys)RI-PSMα3 peptide, the thermodynamic equilibrium distribution shifted nearly quantitatively towards the hetero-dimeric product (*Figure 4D, Expt. No. 5*).

**Figure 4.**
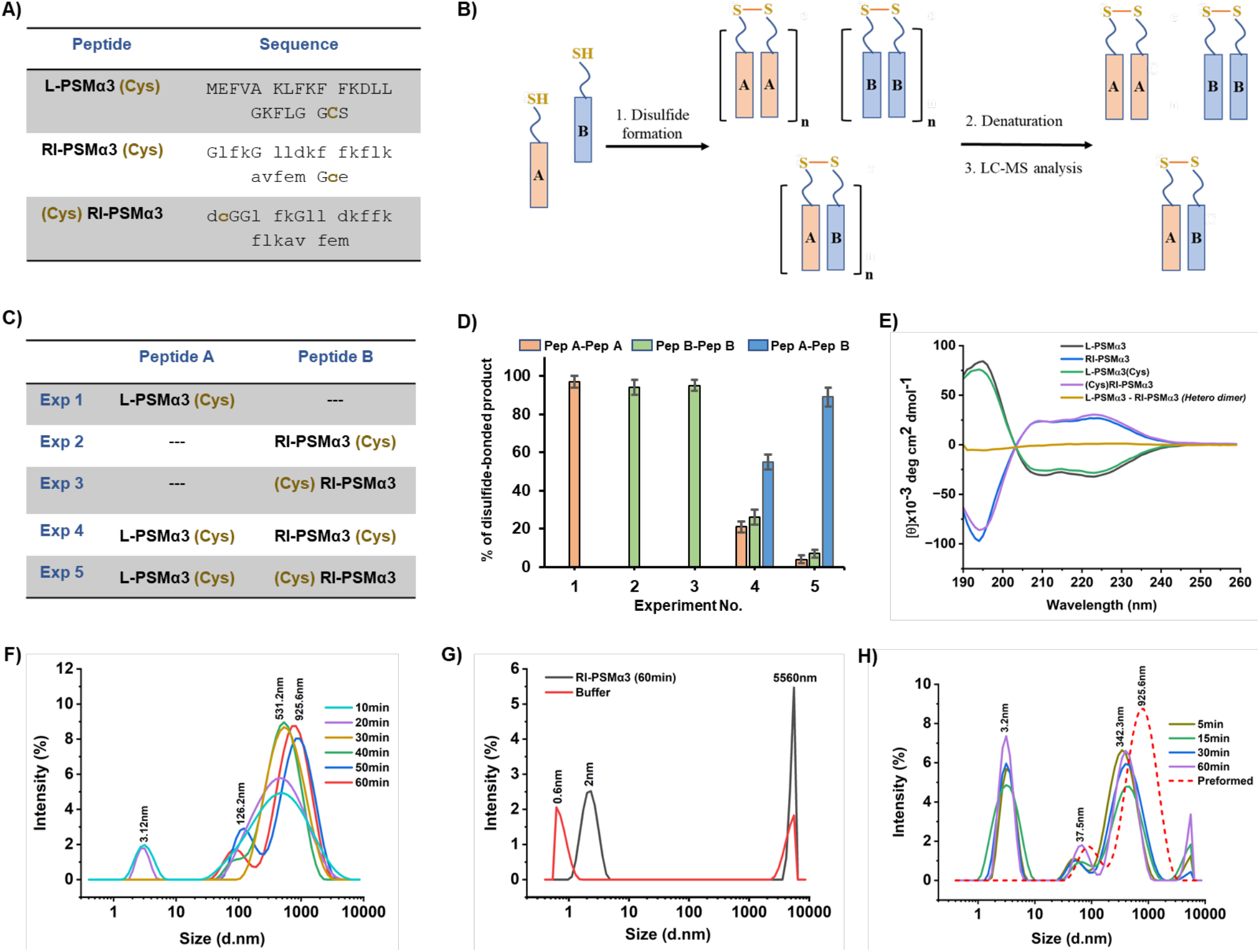
Dynamic disulfide bond formation kinetics, circular-dichroism, and dynamic light scattering experiment. (A) Sequence of the chemically synthesized PSMα3 peptide and its analogs. (B) The schematic representation of two-component dynamic combinatorial disulfide formation of peptides. The orange color bar is the A-type peptide and the blue color bar is the B-type peptide as shown in Table C. The disulfide-bonded complex can be detected after the guanidine-mediated denaturation and RP-HPLC ESI-MS analysis. (C) In the table, the A-type peptide represents the L-PSMα3 peptide having C-terminal cysteine residue while the B-type peptide represents any of the two other peptides having Cys-residue either at the N-terminus or the C-terminus (RI-PSMα3(Cys) and (Cys)RI-PSMα3). (D) An equal equivalent amount of peptide-A was mixed with peptide-B in the folding condition in the presence of redox reagent (20 mM phosphate, 1 mM GSH, 0.2 mM GSSG, pH 8.0). For experiments 1-3, the individual peptide was the only component whereas for experiments 4-5, the mixture of two peptides has been used. The column chart represents the quantity of individual dimer (percentage) obtained when the reaction reached equilibrium. The percentage of the products was quantified from the peak intensity (area under the curve) detected at 214 nm wavelength in analytical RP-HPLC. (E) The far-UV CD spectrum (190nm-260nm) was obtained by measuring the ellipticity of the sample in the aqueous phosphate buffer (20 mM phosphate, pH 7.4). All the peptides showed characteristic alpha-helical signatures with L-peptides showing characteristic negative minima at 208 nm and 222 nm while the RI-peptides showed positive maxima at the same two wavelengths. (F) DLS data of L-PSMα3 aggregating species were taken at different time points and represented as color traces. (G) DLS data of RI-PSMα3 peptide and buffer alone. Only the data point after 1 hour of the incubation of RI-PSMα3 is shown here. (H) DLS data of L-PSMα3 aggregating species taken after 1-hour incubation and the addition of the equivalent amount of the RI-PSM3 peptide. The aggregating species at different time points are represented as color traces.

The dynamic disulfide bond formation chemistry described above revealed two key findings. First, RI-PSMα3 preferentially binds to the L-PSMα3 peptide rather than to itself, indicating a strong interaction between the two peptides when properly oriented. Second, their binding mode favors an antiparallel orientation, providing important insights into the structural aspects of this interaction.

### L-PSMα3 and RI-PSMα3 interact with each other through alpha-helices

The secondary structure of the L-PSMα3 and RI-PSMα3 peptides in the buffer (20 mM phosphate, pH 7.4) was determined individually by recording far-UV circular dichroism (CD) spectra (*see SI Section 1.11*). The CD spectra revealed alpha-helical nature of both peptides, with the RI-PSMα3 peptide showing the reverse CD signature, as expected for a D-peptide (*Figure 4E*). To quantify their equilibrium secondary structure components in the buffer, we used a server-based algorithm BESTsel.^59^ The results indicated that both peptides displayed an equal proportion of alpha-helical content (*See SI Table S2*).

The consistency in the secondary structure content across peptides, as observed from the far-UV CD spectra, supports the notion that PSMα3 and RI-PSMα3 interact with each other through alpha-helices during dynamic disulfide bond formation. Moreover, the straight-line pattern in the CD signature (*Figure 4E*) of the purified hetero dimer product (from experiment No. 5, Figure 4C) further confirms the alpha-helical interaction interface between the two peptides.

### RI-PSMα3 captures transient oligomeric species of L-PSMα3

To gain deeper insights on the underlying mechanism, we conducted dynamic light scattering (DLS) experiments with L-PSMα3 and RI-PSMα3 and explored their fascinating interplay (*see SI Section 1.12*). In the first experiment, L-PSMα3 and RI-PSMα3 peptides were allowed to evolve independently to an equilibrium characterized by a maximally soluble higher oligomeric state. DLS data collection at 10-minute intervals painted a revealing picture over an hour. We found that the population distribution of the L-PSMα3 peptide reached equilibrium, with significant enrichment of species exhibiting average hydrodynamic radii of 925.6 nm (*Figure 4F*). This equilibrium coexisted with minor species having hydrodynamic radii of 3.12 nm and 126.2 nm, indicating their participation in the complex peptide ensemble. In stark contrast, the population distribution of RI-PSM3 remained skewed toward species with very low hydrodynamic radii, residing primarily at 2.0 nm. In the subsequent experiment, we observed a significant shift in the DLS profile when an equal proportion of RI-PSMα3 was added to the pre-equilibrated PSMα3 oligomeric species. The dynamics of the population distribution were monitored at four different time points within an hour (5 minutes, 15 minutes, 30 minutes, 60 minutes). A remarkable finding was revealed during the experiment. The initially dominant population of species with hydrodynamic radii of 925.6nm underwent a profound shift, transitioning to significantly lower-order oligomers characterized by hydrodynamic radii of 342.3nm. At the same time, equilibrium was maintained with the remaining smaller species at 3.2nm, with both populations exhibiting nearly equal intensity (*Figure 4H*).

These careful experiments revealed an intriguing facet of RI-PSMα3 function. We believe that the RI-PSMα3 possesses a remarkable ability to capture transient oligomeric species along the intricate cross-alpha amyloid formation pathway. By doing so, it drives the equilibrium toward specific lowly-populated oligomeric states, suggesting a dynamic and versatile role for RI-PSMα3 in modulating amyloid formation.

### RI-PSMα3 inhibits MRSA biofilm in a dose-dependent manner

Infections caused by *S. aureus* biofilms exhibit significant resistance to antibiotics and the host immune system, making eradication efforts more complex and less effective. This increased resistance complicates treatment strategies and often requires more aggressive or longer-term approaches to treat and eliminate these persistent infections. We embarked on an experiment aimed at dismantling the matured and inhibiting the formation of the EPS matrix of *S. aureus*. Remarkably, a substantial reduction in the biofilm mass in the bacterial population (MRSA strain ATCC 43300) was observed in a crystal-violet staining assay when RI-PSMα3 peptide was applied to a matured biofilm-containing bacterial colony in a dose-dependent manner *(Figure 5A)*. The RI-PSMα3 at 60 µM led to ∼50% inhibition in the biofilm biomass formation. This investigation serves as a compelling demonstration of the singular anti-biofilm potential exhibited by the RI-PSMα3 peptide. Furthermore, the SEM analysis of MRSA biofilm revealed the inhibitory effect of RI-PSMα3 on *S. aureus* biofilm formation. Under the treatment of RI-PSMα3, we observed fewer bacterial cells and less extra-polymeric substance (EPS) accumulation on glass coverslips (*Figure 5B*), which was in agreement with the results obtained from the crystal violet spectrophotometry assay.

**Figure 5.**
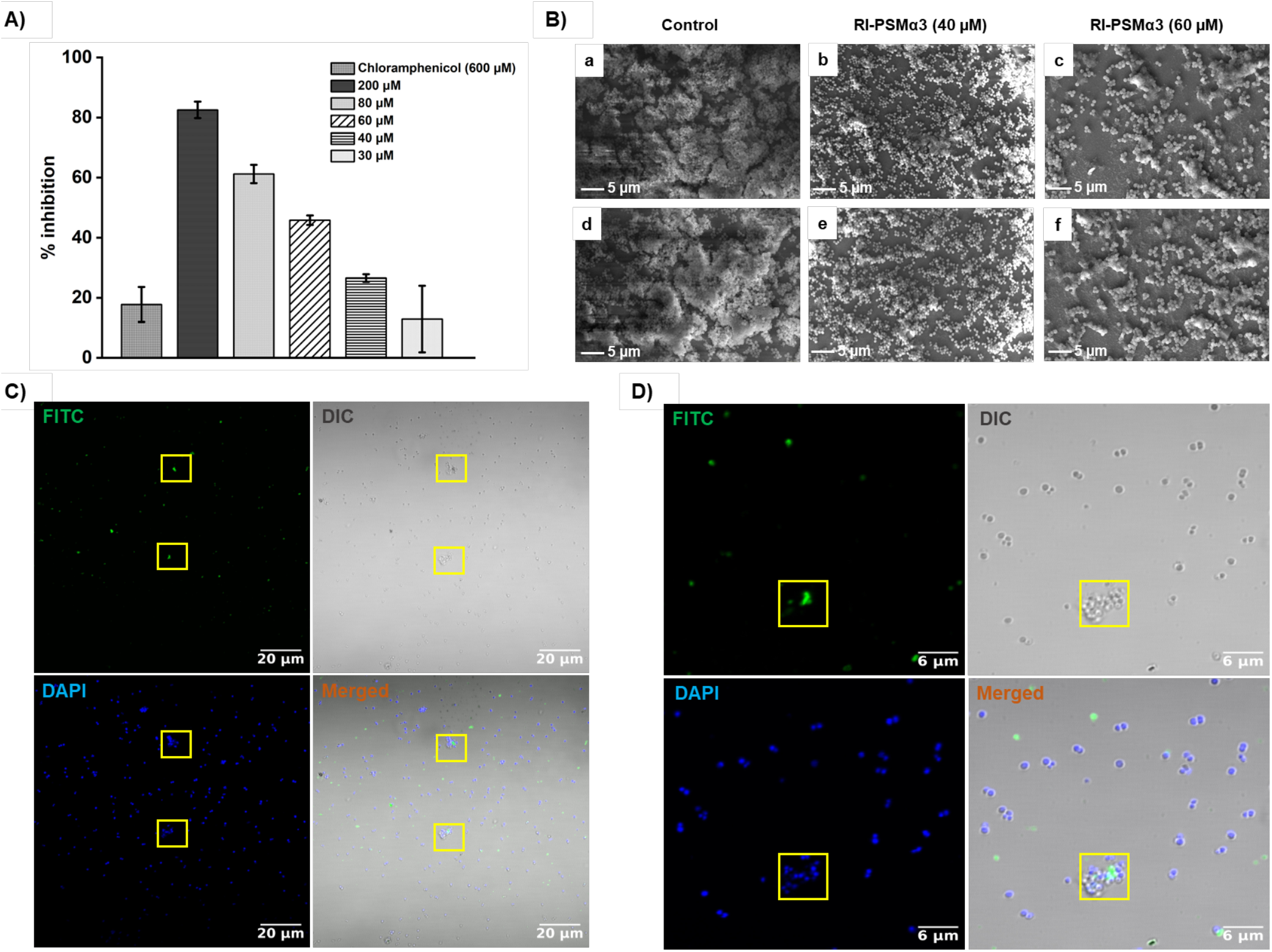
Dose-dependent MRSA biofilm inhibition activity of RI-PSMα3. (A) Percentage MRSA biofilm inhibition by RI-PSMα3. Chloramphenicol was used as a control. B) Micrographs of accumulated biofilms of the MRSA strain (ATCC 43300) formed on glass surfaces upon pre-treatment with RI-PSMα3 at different concentrations as observed by scanning electron microscopy (Carl Zeiss, Evo 18). (a, d) Growth control; (b, e) RI-PSMα3 at 40 µM; and (c, f) RI-PSMα3 at 60 µM. C), D) Colocalization of FITC-tagged RI-PSMα3 with MRSA biofilm. Cells were treated with FITC-tagged RI-PSMα3 and stained with DAPI and visualized under 63X magnification of Leica, TCS SP8 Advanced Spectral Laser Scanning Confocal Microscope. Scale bar = 20 μm (C); Scale bar = 6 μm (D).

### Colocalization of FITC-Tagged RI-PSMα3 Peptide with MRSA Biofilm Matrix

The colocalization study showed that the RI-PSMα3 peptide labeled with FITC got localized with the MRSA biofilm matrix. As shown in the figure (Figure 5C and 5D), a strong fluorescence signal was observed in the biofilm indicating that the labeled peptide had successfully penetrated and attached to the components of the biofilm. The overlap of FITC signals with biofilm structures confirms that RI-PSMα3 interacts effectively with biofilm. This co-location supports the proposed mechanism of action, where RI-PSM3 directly interferes with the integrity of the biofilm by directly binding to its amyloid components. These findings further confirm the potential of RI-PSM3 peptide as a promising agent for the direct targeting and dispersal of infections associated with the biofilm.

## Conclusions

The retro-inverso PSMα3 (RI-PSMα3) peptide, described in this work, holds tremendous potential to function as an effective inhibitor of cross α-helical amyloid formation of native PSMα3 peptide. We found that the RI-PSMα3, which does not form fibrils at high concentrations, inhibits PSMα3 amyloid formation in a dose-dependent manner, as shown by ThT assays. SEM analysis has further confirmed this inhibitory activity by examining amyloid formation of the PSMα3 in presence of varying concentrations of RI-PSMα3 peptide. Additionally, RI-PSMα3 efficiently disassembles pre-formed amyloid aggregates, as evidenced by a dose dependent reduction in ThT fluorescence. The SEM analysis of matured PSMα3 fibrils in the absence or presence of different equivalents of the RI-PSMα3 peptide corroborates the ThT binding assay results. Dynamic combinatorial disulfide bond formation chemistry has established that the retro-inverso PSMα3 peptide preferentially binds with the native PSMα3 in an antiparallel fashion. The dynamic light scattering experiment reveals that the RI-PSMα3 peptide traps transient oligomeric species along the cross-alpha amyloid forming pathway of PSMα3, driving the equilibrium toward specific lowly-populated oligomeric states and providing key mechanistic insight. RI-PSMα3 disperses MRSA biofilm biomass in a dose-dependent manner. Colocalization analysis further shows that FITC-labeled RI-PSMα3 binds to the biofilm matrix, supporting its potential as a direct biofilm-targeting and dispersal agent.

The above findings suggest that RI-PSMα3 could serve as a potential therapeutic agent, advancing our understanding and revolutionizing treatment options for biofilm-associated MRSA bacterial infections. We believe that the dispersal of a biofilm by RI-PSMα3 peptide, coupled with other biofilm-dispersing small molecules and antibiotic therapy, can synergistically eliminate *S. aureus* infection, which is otherwise notoriously difficult to treat.

## Acknowledgments

This research was supported by the intramural funds at TIFR Hyderabad from the Department of Atomic Energy (DAE), Government of India, under Project Identification No. RTI-4007. The authors acknowledge the instrumental facility of the Tata Institute of Fundamental Research, Hyderabad and BRIC-National Institute of Animal Biotechnology (NIAB), Hyderabad AB was supported by SERB Grant No: CRG/2021/003634.

## Entry for the Table of Contents

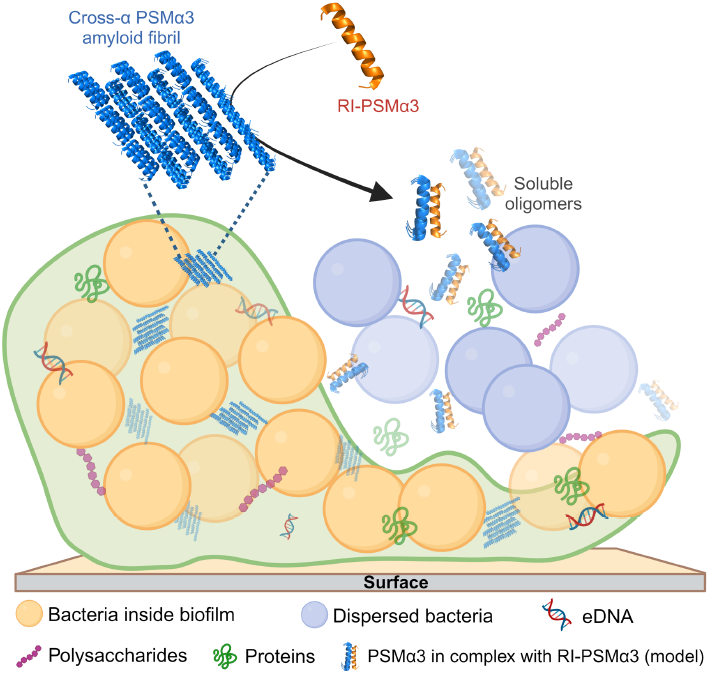

