## Supporting Information for "A Retro-Inverso Peptide that Inhibits Formation and Disrupts Preformed α-Helical Amyloid and Staphylococcus aureus Biofilms"

### 1.1 Chemicals

N,N-diisopropylethylamine (DIEA), Ethyl cyanohydroxyiminoacetate (Oxyma) and all the L- and D-amino acids for Fmoc-SPPS was obtained from Chem-Impex International, USA and Gyros Protein Technologies. The side-chain protecting groups used were, Glu(OtBu), Asp(OtBu), Asn(Trt), Lys(OtBu), Ser(tBu) and Cys(Trt). Dichloromethane (DCM), diethyl ether, N,N'-dimethylformamide (DMF), HPLC grade N,N'-diisopropylcarbodiimide (DIC), 1,1,1,3,3,3-Hexafluoroisopropanol (HFIP), 2,2,2 – Trifluoroethanol (TFE), phenol, 2,2 -(Ethylenedioxy)diethanethiol (DODT), Triisopropylsilane (TIPS) and trifluoroacetic acid (TFA) were purchased from SRL chemicals India. The HPLC grade acetonitrile ($CH_3CN$) for the peptide purification was purchased from Thermofisher Scientific, India. Piperidine was obtained from AVRA chemicals, India. Thioflavin T (stain for amyloid grade) was obtained from Sigma-Aldrich. The Rink amide aminomethyl resin (polystyrene resin with 1% cross-linked with divinylbenzene) was obtained from Supra Science Private Limited, India.

### 1.2 Peptide synthesis, purification, and mass analysis

All peptides (see *Table S1*) were synthesized using a peptide synthesizer (Tribute-UV/IR from Protein Technologies, USA). Fmoc-SPPS was carried out following the reported[60] protocol with minor modifications, using amino acids (AA) (0.25 M), DIC (0.25 M) as a coupling reagent, and oxyma (0.25 M) with DIEA (0.025 M) as additives. Peptides were synthesized on Rink amide aminomethyl resin with a loading capacity of 0.5-0.6 mmol/g. Cysteine was coupled for 5 min at room temperature followed by 10 min at 50 °C and all other amino-acid coupling was performed for 7 min at 65 °C under an $N_2$ atmosphere with vortex mixing. Fmoc deprotection after every coupling cycle was carried out by 20% piperidine in DMF at 50 °C. After synthesis, a cocktail of TFA (85%), phenol (5%), water (5%), DODT (2.5%), and TIPS (2.5%) was used to cleave peptides from the resin. TFA was evaporated to reduce the solution volume, the cleaved peptide was precipitated with diethyl ether, and lyophilized. Analytical reverse-phase high-performance liquid chromatography (RP-HPLC) was performed (*see Figure S1*) to assess the purity of cleaved peptides on an Agilent HPLC instrument using an Agilent zorbax SB-C3 (5 μm), 4.6×150 mm reverse-phase silica column at a flow rate of 0.9 mL/min using a linear gradient of 15-75% solvent B (0.08% TFA in acetonitrile) in solvent A (0.1% TFA in H2O) in 10 minutes. The UV absorbance of the column eluent was monitored at 214 nm wavelength. Preparative (RP-HPLC) of crude peptides was performed on a Waters 1525 preparative HPLC system using Agilent ZORBAX-SB C3 (5 μm, 80 Å, 9.4 × 250 mm) columns at 40 °C using a linear gradient of 25-55% solvent B (0.08% TFA in acetonitrile) in solvent A (0.1% TFA in $H_2O$) in 60 minutes at 45 °C. Fractions containing the purified target peptide were identified by ESI-MS (*see Table S1)*. The deconvolution of the observed mass was carried out using Agilent MassHunter Qualitative Analysis software (version B.07.00), and the deconvoluted mass was reported with an uncertainty of ± 0.02 Da.

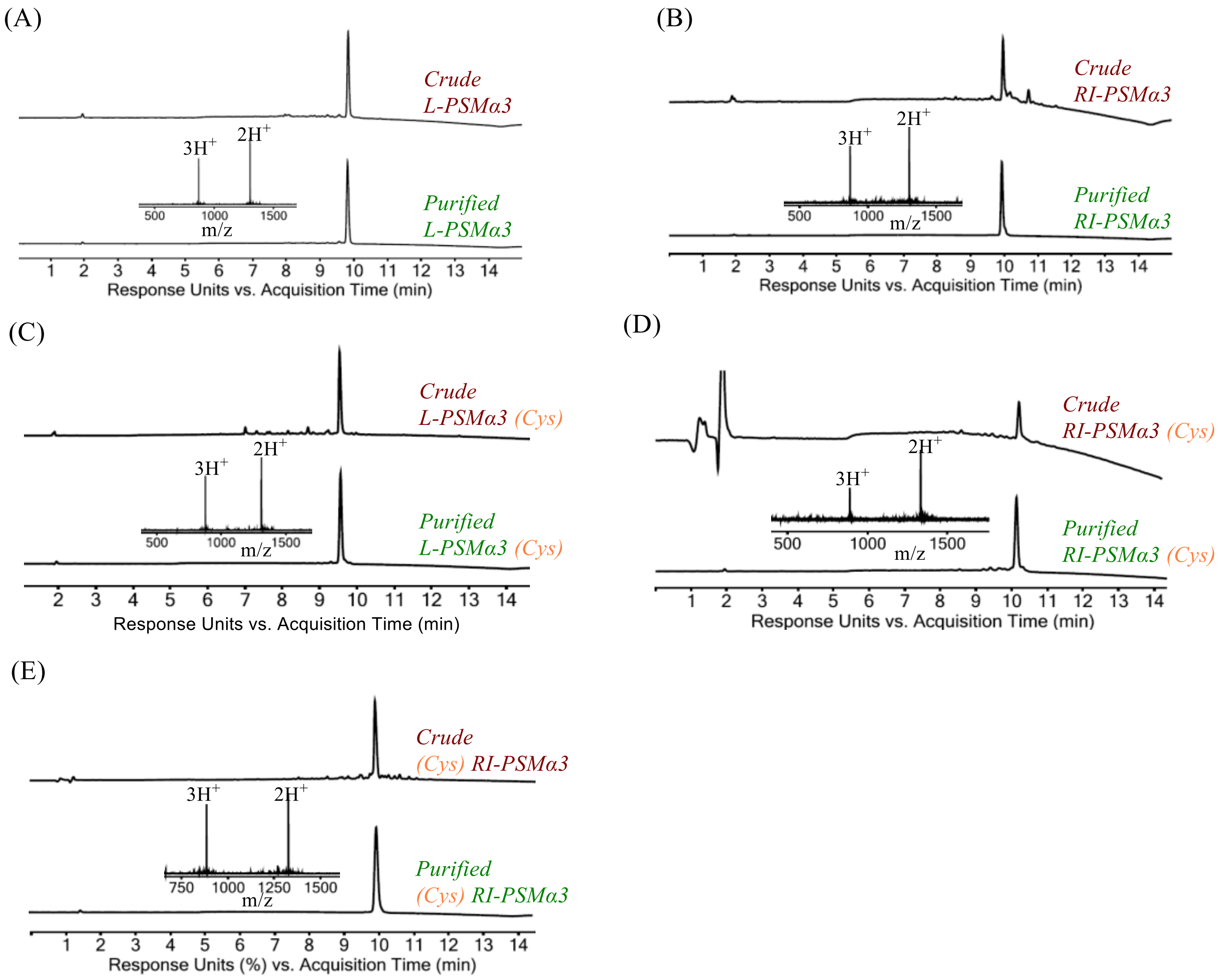


***Figure S1.*** *Analytical RP-HPLC profile (λ = 214 nm) together with ESI-MS data (inset) of the crude and purified peptides. A linear gradient of 15%-75% buffer B in buffer A (buffer A = 0.1% TFA in water; buffer B = 0.08% TFA in acetonitrile) at 40 °C over 10 minutes including 4 min equilibration time using Agilent Zorbax SB-C3, 5 µm, 4.6 x 150 mm, LC column with 0.9 mL/min flow rate was used for the chromatographic separation.*

***Table S1.*** *Sequence of the wild type PSMα3 peptide and the synthesized analogs.*

| Peptide | Sequence** | Purified yield (%) | Calculated mass | Observed mass |
|---|---|---|---|---|
| L-PSMα3 | MEFVAKLFKF FKDLLGKFLG NN | 42 | 2605.41 Da. | 2605.43 Da. |
| RI-PSMα3 | nnGlfkGlld kffkflkavf em | 49 | 2605.41 Da. | 2605.43 Da. |
| L-PSMα3 (Cys) | MEFVAKLFKF FKDLLGKFLG GCS | 55 | 2624.39 Da. | 2624.38 Da. |
| RI-PSMα3 (Cys) | GlfkGlldkf fkflkavfem Gce | 48 | 2666.40 Da. | 2666.39 Da. |
| (Cys) RI-PSMα3 | dcGGlfkGll dkffkflkav fem | 53 | 2652.38 Da. | 2652.40 Da. |

***The capital letters denote L-amino acids and the small letters denote D-amino acids.*

### 1.3 Peptide pretreatment

The lyophilized powder of the L-PSMα3 and RI-PSMα3 peptide analogs were reconstituted in a mixture of TFA-HFIP (1:1) to achieve a concentration of 2 mg/ml. Next, the solution was subjected to sonication for a duration of 10 minutes at a temperature of 37°C. The solution was then allowed to evaporate in a chemical hood over a period of 2 days. To remove any remaining solvent, a high vacuum apparatus was used for solvent evaporation. In cases where immediate testing was not performed, the treated peptides were stored at a temperature of -80°C.

### 1.4 General protocol for stock peptide solution preparation

One portion of the pretreated purified fraction of peptide was dissolved in ultra-pure milli-Q water in an ice bath and sonicated for 10 minutes. The resulting stock mixture was diluted to achieve a final concentration of 1 mM. The concentration was measured using IMPLEN Spectrophotometer (NanoPhotometer® NP80) from Beer-Lambert equation at 205 nm: $A205 = \varepsilon205 \times l \times C$, where l = pathlength in cm, A205 = absorbance value at 205 nm, ε205 = sequence specific extinction coefficient obtained from the web-server (https://bestsel.elte.hu/extcoeff.php).

### 1.5 Protocol for aggregation kinetics study of peptides using Thioflavin-T (ThT) dye

The ThT fluorescence assay protocol was adopted with minor modifications from the protocol described in recent articles.[38] 5 µL of the peptide stock (1 mM, see *Section 1.4*) was mixed with 40 µL of the freshly prepared solution of ThT (1 mM) in the same phosphate buffer saline (10 mM phosphate and 150 mM NaCl, pH 7.4) and 155 µL phosphate buffer saline in 96-well Microplates (Molecular Probes® 96-well microplates). The final concentration of the peptide was 25 µM and ThT was 200 µM. The 96-well plate was immediately covered with silicon sealing film and incubated in a plate reader (POLARstar® Omega) at 37 °C with 400 rpm constant shaking for 30 seconds before each reading collection. The ThT fluorescence ($\lambda_{ex}/\lambda_{em}$ = 438 ± 20 nm/480 ± 20 nm) data was collected for 14 hours with 5 minutes interval each. All experiments were performed in triplicates, subtracted with the blank, and normalized with the maximum signal obtained. Also, standard errors were calculated and represented as error bars in the plots. The entire experiment was repeated at least three times for better reproducibility. The aggregation kinetics of the RI-PSMα3 peptide was subsequently screened for increased concentrations of 50 µM, 100 µM, and 200 µM following the same protocol as above, only by varying the volume of the stock peptide against phosphate buffer saline keeping the ThT concentration in all the cases same.

#### 1.6 General protocol of the scanning electron microscopy (SEM) analysis of fibril

Fibril samples were diluted to 50-fold in ultra-pure water and vortexed gently. The sample solution (5 µL) was then deposited onto clean silicon coverslips, which had been previously tested for effective sample adherence and kept for 10 minutes. Then the sample was washed gently by pouring 10 µL of ultra-pure water and carefully taking out the solvent by micropipette. This process was repeated 5 times and then the sample on silicon slip was allowed to air-dry under controlled conditions to minimize sample disruption overnight. Control samples, such as clean glass coverslips or buffer alone, were also prepared and imaged under the same conditions to account for background and instrument variations. The scanning electron microscopy (SEM) images of amyloid fibrils were captured using an SEM system (JEOL JSM-7200F) with an acceleration voltage of 30 kV. The acquired SEM images were subjected to post-processing. The entire SEM imaging process was repeated multiple times to ensure the reproducibility and consistency of the observed fibrillar appearance.

#### 1.7 Cross-alpha amyloid formation of L-PSMα3 peptide in presence of the RI-PSMα3 peptide

To assess the influence of the RI-PSMα3 peptide on the aggregation kinetics of L-PSMα3, an investigation was undertaken by mixing the equivalent amount of both peptides by varying the concentration of each peptide. The initial step involved preparing 1 mM stock solutions for L-PSMα3 and RI-PSMα3 peptides using ultra-pure water, following the detailed protocol provided in *Section 1.4*. Subsequently, 2.5 µL of each peptide was meticulously combined from the stock solutions. The peptide was then mixed with 40 µL of freshly prepared 1.0 mM ThT solution in PBS (pH 7.4). A final volume of 200 µL was achieved by adding 155 µL of PBS resulting in a final peptide concentration of 12.5 µM, while the ThT concentration remained constant at 200 µM. Data acquisition spanned a period of 14 hours, aligning with the protocol outlined in *Section 1.5*. During this period, changes in ThT fluorescence were recorded to analyze the aggregation kinetics. Similarly, the impact of RI-PSMα3 peptide on L-PSMα3 aggregation kinetics was explored at concentrations of 25 μM and 50 μM for each peptide. The entire experimental procedure was replicated a minimum of three times to ensure reproducibility and consistency of results.

#### 1.8 Dose-dependent effect of RI-PSMα3 peptide on L-PSMα3 peptide aggregation kinetics

A systematic study was conducted to investigate the concentration-dependent influence of RI-PSMα3 peptide on the aggregation kinetics of L-PSMα3. Different concentrations of RI-PSMα3 peptide were combined with L-PSMα3 peptide to gauge their effects on aggregation behavior. Stock solutions of both L-PSMα3 and RI-PSMα3 peptides were meticulously prepared using ultra-pure water in accordance with the procedure outlined in *Section 1.4*. While the concentration of L-PSMα3 peptide remained constant at 25 μM, the concentration of RI-PSMα3 peptide was systematically varied across a range of concentrations: 1.25 μM, 2.50 μM, 3.75 μM, 6.25 μM, 9.25 μM, 12.50 μM, 18.75 μM, and 25.00 μM. For each experimental run, the peptides were combined with 40 μL of freshly prepared 1.0 mM ThT solution in phosphate-buffered saline (PBS; 10 mM phosphate and 150 mM NaCl, pH 7.4). The mixture was then adjusted with an appropriate volume of PBS to achieve the specified peptide concentrations. These preparations were performed in Molecular Probes® 96-well microplates, resulting in a final volume of 200 μL with ThT concentration maintained at 200 μM. Data acquisition was conducted over a 14-hour period, in line with the procedure detailed in *Section 1.5*. This involved monitoring changes in the ThT fluorescence signal to reflect the aggregation kinetics. To ensure the robustness of the results, all experiments were conducted in triplicate and repeated three times for reproducibility.

#### 1.9 Dose-dependent disaggregation of L-PSMα3 peptide with RI-PSMα3 peptide

To investigate the potential of the RI-PSMα3 peptide to destabilize fibrillar structures formed by L-PSMα3, a series of experiments were conducted. Pre-formed L-PSMα3 fibrils were subjected to varying concentrations of RI-PSMα3 peptide, and the resulting reduction in ThT fluorescence signal was used for quantitative assessment of fibril disruption. Initially, L-PSMα3 peptide aggregates were allowed to form over a period of 4 hours in the absence of RI-PSMα3 peptide. This aggregation process was conducted following the procedure outlined in *Section 1.5*. The final concentration of L-PSMα3 peptide in the aggregation mixture was 25 μM, and ThT was added at a concentration of 200 μM in a total volume of 200 μL. The selected duration of 4 hours was determined based on prior experimentation, during which ThT fluorescence intensity reached a stable plateau, indicating complete fibril formation. Subsequently, different quantities of RI-PSMα3 peptide (6.25 μM, 12.50 μM, 18.75 μM, 25.00 μM, 37.50 μM, and 50.00 μM) were introduced to the pre-formed L-PSMα3 fibrils. The RI-PSMα3 peptide was sourced from a 1 mM stock solution, as detailed in *Section 1.4*. The fibril-peptide mixtures were then subjected to an incubation period of 14 hours. Data collection commenced immediately after the addition of the RI-PSMα3 peptide, maintaining a temperature of 37 °C with continuous shaking at 400 rpm for 30 seconds prior to each reading. Readings were taken at 5-minute intervals throughout the incubation period. All experiments were performed in triplicate, and appropriate blank subtraction was applied to the obtained data. Normalization was carried out using the maximum signal acquired for each individual experiment.

#### 1.10 Dynamic disulfide bond formation and reaction monitoring by RP-HPLC and ESI-MS

The cysteine-containing peptides were mixed based on the experiment number as mentioned in *Figure 4C,* under folding condition. The folding buffer consisted of 20.0 mM phosphate, 0.2 mM oxidized glutathione, and 1.0 mM reduced glutathione at pH 8.00. The reversible exchanges of disulfide bonds occur among the thiol groups of cysteines, leading to the formation of various disulfide-linked molecules. This dynamic equilibrium allows the constant interconversion of hetero and homo molecular species. Based on the thermodynamic stability, the population of some species will be higher than the others. The first three experiments (*see table of Figure 4C (exp 1-exp 3)*) were to check individual peptide's ability to form disulfide bond under folding condition. In the next two experiments, the peptides were mixed pairwise as shown in the table of Figure 4C (*exp 4 and exp 5*), maintaining the equivalent amount of each peptide. The final concentration of each peptide was 25 μM.

#### 1.11 Secondary structure analysis of the peptides by circular dichroism spectroscopy

The circular dichroism spectra of the L-PSMα3, RI-PSMα3, L-PSMα3-Cys, and Cys-RI-PSMα3 were measured to evaluate the secondary structure content. A 25 μM solution of peptide in phosphate buffer (20 mM phosphate, pH=7.4) was prepared from the stock solution (1 mM, *see Section 1.4*) to obtain the spectrum for monomeric PSMα3. Spectra for each peptide was recorded using a Jasco J-1500 CD spectrometer at room temperature by scanning wavelength from 260 nm to 190 nm. The measurements represent the average of three scans and were subtracted with the appropriate blank.

Mean residual ellipticity was calculated using the following equation,

Mean residual ellipticity (deg.cm2.dmol-1) = (CD value (mDeg) * Mw) / (10*c*l*Nr)

Where Mw is the molecular weight of the protein, c is concentration in g/L, l is the path length of the cuvette in cm and Nr is the number of amino acid residues in the peptide. The concentrations of all peptides were measured using IMPLEN Spectrophotometer (NanoPhotometer® NP80) from the Beer-Lambert equation as discussed in *Section 1.4.*

We used a server-based computational algorithm (BESTsel)[24] for quantitative analysis of the secondary structure content. The percentage of helicity calculated is shown in *Table S2*.

***Table S2.*** *Percentage of helicity calculated using BESTsel.*

| Peptide | % of helicity |
|---|---|
| L-PSMα3 | 52.8 |
| RI-PSMα3 | 40.8 |
| L-PSMα3 (Cys) | 45.2 |
| RI-PSMα3 (Cys) | 51.3 |
| (Cys) RI-PSMα3 | 52.4 |

### 1.12 Size distribution analysis of L-PSMα3 peptide in the presence and absence of the RI-PSMα3 peptide

Dynamic Light Scattering measurements were conducted at 25 °C using a Malvern Zetasizer Nano ZS (Malvern Instruments, UK). The instrument was equipped with a 10 mW helium-neon laser operating at a wavelength of 632.8 nm and a scattering angle set at 173°. Each measurement was carried out with correlation times defined over a 10-second duration per run, and a total of 20 runs were performed for each measurement. Samples were prepared in a 10 mM phosphate buffer with 150 mM NaCl at pH 7.4. Prior to measurement, samples were subjected to ultra-centrifugation and filtration through 0.2 µm filters. Subsequently, the prepared samples were transferred to the DLS cuvette for analysis. Data collection was carried out at 10-minute intervals over the course of 1 hour. The resulting data was plotted to depict the distribution intensity (%) of particles relative to their hydrodynamic radius (nm). The experimental process began by examining the distribution pattern changes for both L-PSMα3 and RI-PSMα3 peptides independently. Subsequently, an additional experiment was conducted, wherein the L-PSMα3 peptide was first allowed to reach to a stable higher-order soluble oligomeric state. At this point, an equivalent amount of the RI-PSMα3 peptide was introduced from the stock solution. The alteration in population size distribution was once again tracked by collecting data at 10-minute intervals over the course of 1 hour. The obtained experimental data were subjected to analysis using the Zetasizer software version 5. 23 provided with the equipment.

#### 1.13 Dose-dependent MRSA biofilm inhibition assay

**Bacteria growth conditions**

Antibiofilm activity of PSMα3 was carried out against standard pathogenic bacterial strain, MRSA (Methicillin-resistant *S. aureus* ATCC 43300) procured from the American Type Culture Collection (ATCC), and maintained on Tryptic soy agar (TSA), (Himedia). Bacterial cultures were obtained by transferring a single colony to a fresh sterile medium and allowed to grow overnight at 37°C, 200 rpm. The turbidity of the culture was adjusted with a sterile medium to achieve a total of 0.5 Mc Farland standard ($10^8$ colony-forming units/mL).

**Assessment of the adherence of Biofilms**

The effect of RI-PSMα3 on the inhibition of biofilm formation in 96-well microtiter plates was accomplished by a spectrophotometric method as described previously.[61] Briefly, 200 µM of bacterial cell suspensions ($10^8$ CFU/mL) and different concentrations of RI-PSMα3 (30 µM, 40 µM, 60 µM, 80 µM, and 200 µM) were incubated at 37°C for 24 h under stationary conditions. Following the incubation period, the planktonic cells were removed from the wells by carefully washing them with sterile 1X PBS. Biofilms developed by adherent cells were stained with 0.1% crystal violet followed by incubation at 37°C for 30 min. The excess stain was washed off with PBS and stained cells were resuspended in 200 µL of 95% ethanol by further incubation at 37°C for 15 min. Absorbance was read spectrophotometrically at 570 nm. The percentage inhibition was estimated:

[OD (control) – OD (test)/OD (control)] X 100

**Biofilm SEM analysis**

*Staphylococcus aureus* ATCC 43300 were seeded in trypticase soy agar (TSA), incubated for 24 h at 37°C, and inoculated into a tube containing 5 mL of Tryptic soy broth (TSB) with 1% glucose. Then, 500 µL of the inoculated broth ($10^8$ CFU/mL) was added to a 12-well plate containing glass coverslips (18 mm). RI-PSMα3 was added to wells with final concentrations of 40 µM and 60 µM. Biofilms on glass coverslips were cultured for 24 h at 37°C. Fixation of cells was done in 2.5% glutaraldehyde in 0.1 M cacodylate buffer (pH 7.4) at 37°C for 30 mins followed by overnight incubation at 4°C. Cells adhered to the coverslip were dehydrated through a graded series of ethanol solutions (20%, 30%, 50%, 70%,

90% for 15 min, and 95% for 1 hr) at each concentration. Cells were then airdried for 30 mins and coverslips were mounted on aluminum holders, sputtered with gold, and analyzed in a scanning electron microscope (Carl Zeiss, Evo 18) for observation of the biofilms and bacterial morphology.

#### 1.14 Confocal Microscopy

Biofilms were allowed to form on glass coverslips (*as discussed in the previous section*) for 6 hr at 37°C. The glass coverslips were washed thrice with sterile 1X PBS, and the cells were treated with sub-inhibitory concentration of FITC-tagged RI-PSMα3 (60 µM) in TSB and further incubated at 37°C for 1 hr. Cells were washed with sterile 1X PBS to remove unbound FITC-tagged RI-PSMα3 and labelled with DAPI for 5 min and observed with dual-channel scanning at 495 nm and 405 nm under Super Resolution Microscope (Carl Zeiss, Elyra, LSM 880).
